# Intergenic ORFs as elementary structural modules of *de novo* gene birth and protein evolution

**DOI:** 10.1101/2021.04.13.439703

**Authors:** Chris Papadopoulos, Isabelle Callebaut, Jean-Christophe Gelly, Isabelle Hatin, Olivier Namy, Maxime Renard, Olivier Lespinet, Anne Lopes

**Affiliations:** Université Paris-Saclay, CEA, CNRS, Institute for Integrative Biology of the Cell (I2BC), 91198, Gif-sur-Yvette, France; Sorbonne Université, Muséum National d'Histoire Naturelle, UMR CNRS 7590, Institut de Minéralogie, de Physique des Matériaux et de Cosmochimie, IMPMC, 75005, Paris, France; Université de Paris, Biologie Intégrée du Globule Rouge, UMR_S1134, BIGR, INSERM, F-75015, Paris, France; Laboratoire d’Excellence GR-Ex, Paris, France; Institut National de la Transfusion Sanguine, F-75015, Paris, France

**Keywords:** *de novo* genes, noncoding genome, foldability, genome evolution, protein evolution, protein bricks

## Abstract

The noncoding genome plays an important role in *de novo* gene birth and in the emergence of genetic novelty. Nevertheless, how noncoding sequences’ properties could promote the birth of novel genes and shape the evolution and the structural diversity of proteins remains unclear. Therefore, by combining different bioinformatic approaches, we characterized the fold potential diversity of the amino acid sequences encoded by all intergenic ORFs (Open Reading Frames) of *S. cerevisiae* with the aim of (i) exploring whether the large structural diversity observed in proteomes is already present in noncoding sequences, and (ii) estimating the potential of the noncoding genome to produce novel protein bricks that can either give rise to novel genes or be integrated into pre-existing proteins, thus participating in protein structure diversity and evolution. We showed that amino acid sequences encoded by most yeast intergenic ORFs contain the elementary building blocks of protein structures. Moreover, they encompass the large structural diversity of canonical proteins with strikingly the majority predicted as foldable. Then, we investigated the early stages of *de novo* gene birth by identifying intergenic ORFs with a strong translation signal in ribosome profiling experiments and by reconstructing the ancestral sequences of 70 yeast *de novo* genes. This enabled us to highlight sequence and structural factors determining *de novo* gene emergence. Finally, we showed a strong correlation between the fold potential of *de novo* proteins and the one of their ancestral amino acid sequences, reflecting the relationship between the noncoding genome and the protein structure universe.

## Introduction

Many studies give a central role to the noncoding genome in *de novo* gene birth (Ingolia et al. 2011; Tautz and Domazet-Lošo 2011; Carvunis et al. 2012; Slavoff et al. 2013; Prabakaran et al. 2014; Ruiz-Orera et al. 2014; Zhao et al. 2014; Schlötterer 2015; Ruiz-Orera et al. 2018; Vakirlis et al. 2018; Schmitz et al. 2018; Durand et al. 2019; Blevins et al. 2021). Nevertheless, how noncoding sequences can code for a functional product and consequently give rise to novel genes remains unclear. Indeed, function is intimately related to protein structure and more generally to protein structural properties. All proteomes are characterized by a large diversity of structural states. The structural properties of a protein result from its composition in hydrophobic and hydrophilic residues. Highly disordered proteins display a high hydrophilic residue content. Membrane proteins which fold in lipidic environments, but aggregate in solution, are enriched in hydrophobic residues. Finally, foldable proteins are characterized by a subtle equilibrium of hydrophobic and hydrophilic residues (Talmud and Bresler 1944). The latter are arranged together into specific patterns that dictate the formation of the secondary structures and the outcoming fold. However, noncoding sequences display different nucleotide frequencies from coding sequences, resulting in different amino acid compositions. If and how these amino acid compositions can account for the structural states observed in proteomes is a crucial question to understand the relationship, if any, between the noncoding genome and the protein structure universe. So far, different models of *de novo* gene emergence have been proposed (Carvunis et al. 2012; Schlötterer 2015; Wilson and Masel 2011). The “preadaptation” model stipulates that only sequences pre-adapted not to be harmful (i.e. with enough disorder not to be subjected to aggregation), will give rise to gene birth (Wilson and Masel 2011). This model is supported by the observation that young genes and *de novo* protein domains display a higher disorder propensity than old genes (Schmitz et al. 2018; Wilson and Masel 2011; Bitard-Feildel et al. 2015; Ekman and Elofsson 2010). On the contrary, several studies conducted on *S. cerevisiae* indicate that young genes are less prone to disorder (Carvunis et al. 2012; Vakirlis et al. 2018, 2020). Recently, Vakirlis et al. (2020) proposed a TM-first model where the membrane environment provides a safe niche for transmembrane (TM) adaptive emerging peptides which can further evolve toward more soluble peptides. These adaptive peptides have been identified with overexpression which, according to the authors, may not be reached outside the laboratory. Whether such peptides, though beneficial in the experimental conditions, would be produced and would be beneficial in “natural” conditions deserves further investigations.

Overall, all these studies attribute to the fold potential of noncoding ORFs (including the propensities for disorder, folded state, and aggregation) an important role in the emergence of genetic novelty. However, several questions remain open. First, if the sequence and structural properties of *de novo* genes have been largely investigated in specific species, the raw material for *de novo* gene birth and the early stages preceding the fixation of the beneficial ORFs are to be further characterized (Schmitz et al. 2018). Second, if the role of the noncoding genome in *de novo* gene birth has been largely investigated, its role in protein evolution and structural diversity is to be further characterized as well. Indeed, *de novo* domains may emerge from noncoding regions through ORF extension or exonization of introns (Bornberg-Bauer and Alba 2013; Bornberg-Bauer et al. 2015). On the other hand, we can assume that protein-coding genes, whatever their evolutionary history, have had a noncoding ancestral origin (Nielly-Thibault and Landry 2019). Whether the noncoding ORFs which gave rise to novel genes can account for the structural diversity of proteomes or whether this structural diversity evolved from ancestral genes which all displayed similar structural properties (i.e. disordered, foldable or TM-prone) is a crucial question to better understand the role, if any, of noncoding sequences in the protein structure universe.

Therefore, by combining different bioinformatic approaches, we characterized the diversity of the fold potential (i.e. propensity for disorder, folded state or aggregation) encoded in all intergenic ORFs (IGORFs) of *S. cerevisiae* with the aim of (i) exploring whether the large structural diversity observed in proteomes is already present in noncoding sequences, thereby investigating the relationship, if any, between the fold potential of the peptides encoded by IGORFs and the protein structural diversity of proteins, and (ii) estimating the potential of the noncoding genome to produce novel protein bricks, that can either give birth to novel genes or be integrated into pre-existing proteins, thus participating in protein structure evolution and diversity. We then characterized the early stages preceding *de novo* gene emergence with two complementary approaches (i) the systematic reconstruction of the ancestral sequences of 70 *de novo* genes, in order to identify the sequence and structural features of the peptides encoded by IGORFs that indeed gave birth in the past to *de novo* genes and (ii) the identification of IGORFs with a translation signal through ribosome profiling experiments, in order to investigate the sequence and structural properties of the peptides encoded by candidate IGORFs that could give birth to future novel genes. In particular, we performed ribosome profiling experiments on *S. cerevisiae* and used additional ribosome profiling data to precisely analyze their translational behavior. Then we characterized the sequence and structural properties of the peptides encoded by IGORFs that are occasionally translated with a weak translation signal and that may give rise to a novel gene and IGORFs that display a strong translation signature in at least two independent experiments probably reflecting the optimization of their translational activity and the emergence of function.

## Results

We extracted 105041 IGORFs of at least 60 nucleotides in *S. cerevisiae* (see Methods). We probed their fold potential with the Hydrophobic Cluster analysis (HCA) approach (Faure and Callebaut 2013; Bitard-Feildel and Callebaut 2017; Bitard-Feildel et al. 2018; Bitard-Feildel and Callebaut 2018) and compared it with the one of the 6669 CDS of *S. cerevisiae*. HCA highlights from the sole information of a single amino acid sequence, the building blocks of protein folds that constitute signatures of folded domains. They consist of clusters of strong hydrophobic amino acids that have been shown to be associated with regular secondary structures (Bitard-Feildel and Callebaut 2017; Bitard-Feildel et al. 2018; Lamiable et al. 2019) (Fig. S1). These clusters are connected by linkers corresponding to loops or disordered regions. The combination of hydrophobic clusters and linkers in a sequence determines its fold potential. The latter can be appreciated in a quantitative way through the calculation of a foldability score (HCA score) which covers all the structural diversity of proteins.

### IGORFs contain elementary building blocks of proteins

We first investigated the structural and sequence properties of protein encoded by CDS and IGORFs. CDS are longer than IGORFs and contain more HCA clusters (Mann-Whitney U-test, P < 2.2e-16 for both observations) (Figure 1AB). Strikingly, the HCA clusters of CDS and IGORFs display a remarkable similar size of about 11 residues (Mann-Whitney U-test, P = 0.17) (Figure 1C) and 96.9% of IGORFs harbor at least one HCA cluster. This result shows that the elementary building blocks of proteins are widespread in noncoding sequences. In contrast, CDS are enriched in long linkers suggesting that linker sizes have increased during evolution (6.3 and 11.5 residues for IGORFs and CDS respectively, Mann-Whitney U-test, P = 2.6e-11) (Figure 1D). This increase in size might favor protein modularity and flexibility, probably reflecting their important role in protein function optimization and protein structure diversity (Blouin et al. 2004; Espadaler et al. 2006; Tendulkar et al. 2004; Papaleo et al. 2016).

**Figure 1.**
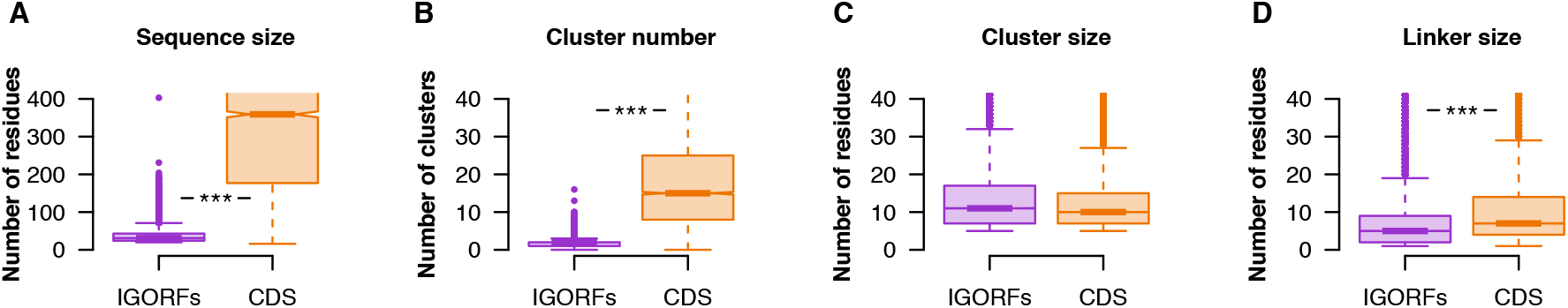
Boxplot distributions of sequence and HCA-based structural properties of IGORFs and CDS (A) sequence size (B) number of HCA clusters per sequence (C) size of HCA clusters (D) size of linkers. Asterisks denote level of significance: ***p < 0.001, see Tables S3-6 for detailed p-values.

### CDS are enriched in polar and charged residues

If hydrophobic clusters of CDS and IGORFs display similar sizes, they may not have the same amino acid composition. Therefore, in order to see whether clusters of CDS have evolved toward specific amino acid distributions, we calculated for each amino acid, its propensity for being in HCA clusters of CDS over HCA clusters of IGORFs. CDS HCA clusters are clearly enriched in polar and charged residues compared to those of IGORFs (Fig. S2A). The same tendency is observed for CDS linkers (Fig. S2B). Strikingly, negatively charged residues are over-represented compared to positively charged ones in both HCA clusters and linkers of CDS (Fig. S2). In fact, it has been shown that the charge distribution in a protein sequence has an impact on its diffusion in the cytosol where positively charged proteins get caught in nonspecific interactions with the abundant negatively charged ribosomes (Requião et al. 2017; Schavemaker et al. 2017). Interestingly, Figure S3 shows that the frequency of negatively charged residues calculated on the yeast cytoplasmic proteins is strongly correlated (Spearman correlation coefficient: Rho = 0.44, P < 2e-16) with the proteins' abundance suggesting that the crowded cellular environment has shaped the charge distribution of abundant proteins. This result recalls the observation made in previous studies which showed that the frequency of “sticky” amino acids on the surface of globular proteins or in disordered proteins decreases as the protein cellular concentration increases (Levy et al. 2012; Macossay-Castillo et al. 2019).

### IGORFs encode for peptides that display a wide diversity of fold potential including a substantial amount of non-harmful peptides

We next used the HCA score in order to assess the fold potential of the peptides encoded by IGORFs. As reference, we calculated the HCA scores for three sequence datasets consisting of 731 disordered regions, 559 globular proteins and 1269 TM regions extracted from transmembrane proteins, thereby expected to form aggregates in solution while being able to fold in lipidic environments (see Methods for more details) (Figure 2A). Based on their HCA scores, we defined three categories of fold potentials (i.e. disorder prone, foldable, or aggregation prone in solution) (Figure 2A). Here, we define as foldable, proteins that are able to fold into a compact and well-defined 3D structure or partially to an ordered structure in which the secondary structures are however present. Figure 2B shows that CDS and IGORFs belonging to the low HCA score category are indeed presumed to be disordered and display low propensity for aggregation. Interestingly, comparable but small proportions of CDS and IGORFs fall into this group (4.9% and 7.7% respectively) indicating that most coding but also noncoding sequences are not highly prone to disorder in line with Tretyachenko et al. (2017). The high HCA score category corresponds to sequences which exhibit a low propensity for disorder while displaying a high propensity for aggregation in solution. CDS falling into this bin correspond to highly hydrophobic sequences (Table S1) among which 81% are annotated as uncharacterized according to Uniprot (UniProt Consortium 2019) and 60% are predicted as containing at least one TM domain. Finally, the intermediate category concerns sequences which have a high potential for being completely or partially folded in solution as shown by their intermediate HCA scores comparable to those of globular proteins. As anticipated, most CDS (91.4%) and, strikingly, a majority of IGORFs (66.6%) fall into this category. Both are characterized by intermediate aggregation and disorder propensities, although IGORFs display a wider range of aggregation propensities (Fig. 2B). The fact that these CDS, though predicted as foldable, exhibit a certain propensity for aggregation, is in line with several studies which report a high aggregation propensity of proteomes across all kingdoms of life (Langenberg et al. 2020; Greenwald and Riek 2012). This observation has been explained as the side effect of the requirement of a hydrophobic core to form globular structures (Langenberg et al. 2020; Ganesan et al. 2016; Rousseau et al. 2006b). In particular, Langenberg et al. (2020), show a strong relationship between protein stability and aggregation propensity with aggregation prone regions mostly buried into the protein and providing stability to the resulting fold. Like for CDS, these regions, under the hydrophobic effect, may facilitate the stabilization of the IGORF encoded peptide structure. Whether peptides encoded by IGORFs in this intermediate category fold to a specific 3D structure, a partially ordered structure or a “rudimentary fold” which stabilizes itself through oligomerization like the Bsc4 *de novo* protein (Bungard et al. 2017), would deserve further investigations. Finally, it is interesting to note that the proportions of sequences in the different fold potential categories are different between IGORFs and CDS, with CDS mostly falling into the intermediate HCA score category reflecting that being foldable is a trait which has been strongly selected by evolution. In contrast, IGORFs cover a wide range of fold potentials. It is questionable whether *de novo* genes mainly originate from IGORFs encoding foldable peptides or from IGORFs whose corresponding peptides subsequently evolved toward foldable peptides regardless of their initial fold potential.

**Figure 2.**
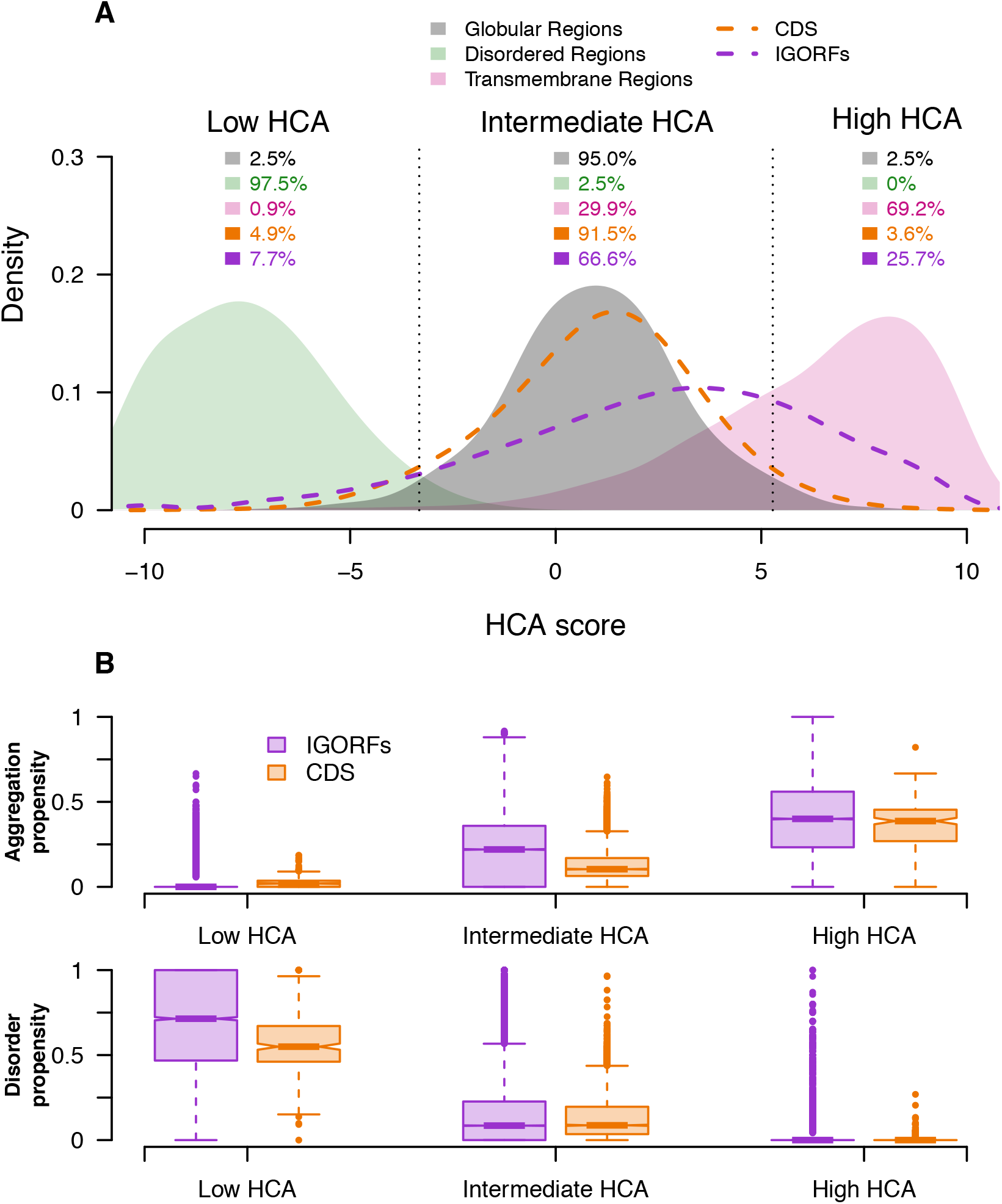
IGORFs encompass the large spectrum of fold potential of canonical proteins (A) Distribution of the HCA scores for the three reference datasets (i.e. disordered regions, globular domains and transmembrane regions - green, black and pink curves respectively) along with those for the CDS (orange curve) and IGORFs (purple curve). There is a clear distinction between the distributions of HCA scores calculated for the three reference datasets. (Two-sided Kolmogorov Smirnov test, P < 2e-16 for all comparisons). Dotted black lines delineate the boundaries of the low, intermediate and high HCA score categories reflecting the three categories of fold potential (i.e. disorder prone, foldable, or aggregation prone in solution). The boundaries are defined so that 95% of globular domains fall into the intermediate HCA score category whereas the low and high HCA score categories include all sequences with HCA values that are lower or higher than those of 97.5% of globular domains respectively. High HCA scores reflect sequences with high densities in HCA clusters that are likely to form aggregates in solution. Low HCA scores indicate sequences with high propensities for disorder, while intermediate scores correspond to globular proteins characterized by an equilibrium of hydrophobic and hydrophilic residues (see Methods for more details). The percentages of sequences in each category are given for all datasets. Raw data distributions are presented in Fig S4. (B) Aggregation and disorder propensities calculated with TANGO and IUPred respectively are given for CDS and IGORFs in each foldability HCA score category.

### From IGORFs to de novo genes

Therefore, we traced back the evolutionary events preceding the emergence of 70 *de novo* genes identified in *S. cerevisiae* by reconstructing their ancestral IGORFs (ancIGORFs) in order to see whether IGORFs that gave birth to *de novo* genes encode peptides that display different foldability potential from all other IGORFs and to characterize the steps preceding the emergence of a novel gene (see Methods and Fig. S5 for more details on the protocol and Table S2 for the list of *de novo* genes). Figure S6 shows the example of YOR333C *de novo* gene which emerged in the lineage of *S. cerevisiae*. The corresponding noncoding region in the ancestors preceding the emergence of YOR333C consists of two IGORFs separated by a STOP codon. Interestingly, two nucleotide substitutions which occurred specifically in the *S. cerevisiae* lineage led respectively to the appearance of a start codon (mutation of Isoleucine into Methionine through an Adenine/Guanine substitution) and the mutation of the STOP codon into a Tyrosine through a Guanine/Cytosine substitution, thereby merging the two consecutive IGORFs. Overall, the 70 *de novo* genes emerged from a total of 167 ancIGORFs. A minority of *de novo* genes (16 cases) emerged from a single ancIGORF which covers almost all their sequence (i.e. single- ancIGORF *de novo* genes), while, the majority (54 cases) result from the combination of multiple ancIGORFs through frameshift events and/or STOP codon mutations as observed with the example of YOR333C (i.e. multiple-ancIGORF *de novo* genes). Interestingly, the multiple-ancIGORF *de novo* genes exhibit sequence sizes similar to those of the single-ancIGORF ones (Fig. S7A) though the ancestral IGORFs they originate from are shorter than those that led to single-ancIGORFs *de novo* genes (Fig. S7B). They evoke the *expression first* model (Schlötterer 2015) where a transcribed and selected IGORF is subsequently combined with neighboring IGORFs through multiple frameshift events and STOP codon mutations. In contrast, single-ancIGORFs *de novo* genes derive from longer ancestral IGORFs and recall the *ORF first* model (Schlötterer 2015) which stipulates that the emergence of a long *de novo* ORF precedes that of the promoter region.

**Figure 3** shows the HCA scores of the proteins encoded by the 70 *de novo* genes (i.e. *de novo* proteins) and of the peptides encoded by their corresponding ancIGORFs. The majority of *de novo* proteins (78%) are predicted as foldable, whereas peptides encoded by ancIGORFs display a larger range of HCA scores. However, ancIGORFs are not IGORF-like, being enriched in sequences encoding foldable peptides (75.4% and 66.6% for ancIGORFs and IGORFs respectively - one proportion z test, P = 9.5e-3) and depleted in sequences encoding aggregation prone ones (18.6% and 25.7% for ancIGORFs and IGORFs respectively, one proportion z test, P = 2.1e-2). This suggests that IGORFs encoding foldable peptide are more likely to give rise to novel genes.

**Figure 3.**
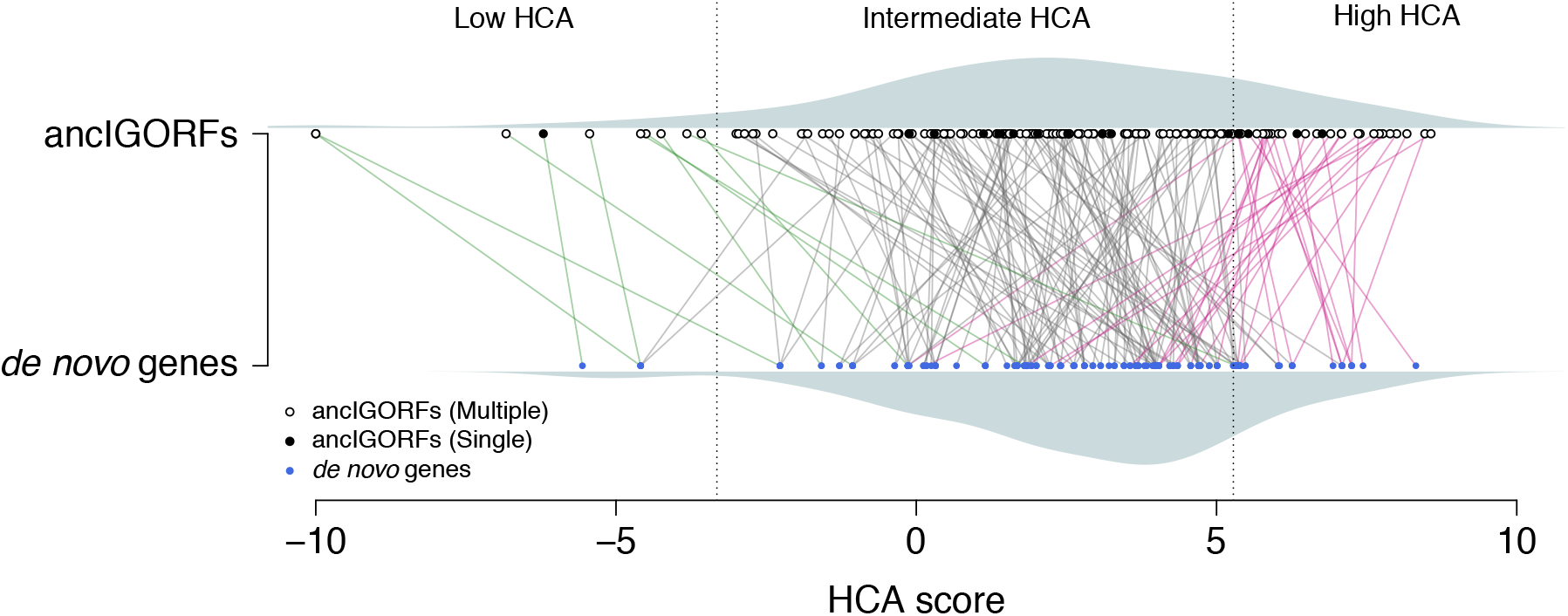
From ancIGORFs to *de novo* genes. Plot of the HCA score of each ancIGORF (black and white points for single and multiple ancIGORFs respectively) along with its corresponding *de novo* gene (blue points). Each *de novo* gene is connected to its parent ancIGORF(s) with a colored line. One should notice that a *de novo* gene can be connected to several IGORFs when it results from the combination of different ancestral IGORFs (i.e. multiple-ancIGORF *de novo* genes). Green lines indicate cases where a *de novo* gene is connected to a low HCA score ancIGORF, while grey and pink lines indicate connections with an intermediate and a high HCA score ancIGORFs, respectively. The HCA score densities of *de novo* genes and ancIGORFs are shown in grey (bottom and top of the graph respectively).

### Impact of frameshift events and STOP codon mutations on the fold potential of a de novo protein

Interestingly, the overall relationship between the HCA scores of peptides encoded by ancIGORFs and their corresponding *de novo* proteins is characterized by a funnel shape revealing that most *de novo* proteins are foldable regardless of the fold potential of the peptides encoded by their IGORF parents (Fig. 3). Two hypotheses can explain this observation: (i) this funnel mostly results from the amino acid substitutions which have occurred since the fixation of the ancIGORF(s) and which led to an increase in foldability of the resulting *de novo* genes, (ii) this funnel results from the fact that combining at least one IGORF encoding for a foldable peptide with IGORFs encoding peptides with different fold potentials, will lead to a foldable product. Figure 4A shows the amino acid frequencies of IGORFs, ancIGORFs, *de novo* genes and CDS. Interestingly, *de novo* genes display amino acid frequencies similar to those of ancIGORFs (blue circles and grey dots respectively) which overall, follow those of all IGORFs (purple line)(see also Table S1 for the frequencies of hydrophobic residues in the different ORF categories). This result shows that the mutations which occurred since the fixation of the ancIGORF did not change the amino acid composition of the resulting *de novo* genes and thus, cannot explain the funnel shape observed in Figure 3. We then reasoned that since the divergence of the last common ancestor predating the emergence of *de novo* genes, single-ancIGORF *de novo* genes were only subjected to nucleotide substitutions, some of which leading to amino acid mutations, while the multiple-ancIGORF ones have undergone frameshift events and/or STOP codon mutations as well. In order to quantify the impact of these different mutational events on the fold potential of the outcoming *de novo* proteins, we calculated the correlation between the HCA score of each *de novo* protein and the peptides encoded by its corresponding ancIGORF(s). Figure 4B shows that single-ancIGORF *de novo* proteins display a clear correlation of HCA scores with those of the peptides encoded by their corresponding ancIGORFs (Pearson correlation coefficient: R = 0.94, P < 2.5e-9). This reveals that the amino acid mutations which occurred between the ancestor and the *de novo* protein did not affect the fold potential of the ancestral sequences. This suggests that the structural properties of the peptides encoded by the single-ancIGORFs were retained in the resulting *de novo* proteins. Interestingly, the correlation is weaker for multiple-ancIGORF *de novo* proteins (Pearson correlation coefficient: R = 0.53, P < 1.6e-7). This can be attributed to the fact that 81% (44/54) of the multiple-ancIGORF *de novo* proteins are predicted as foldable (white dots included in the blue squares in Figure 4B) while being associated with ancIGORFs of different foldability potentials. Interestingly, all foldable *de novo* genes include at least one foldable ancestral peptide suggesting that in these cases, combining disordered or aggregation-prone peptides with a foldable one, has led to a foldable *de novo* protein as well. Figure S5E shows the example of the *de novo* gene YLL020C and its corresponding ancIGORFs. YLL020C results from the combination through a frameshift event, of a long foldable ancIGORF with a short IGORF predicted as aggregation prone. Interestingly, the resulting *de novo* gene is also predicted as foldable. Whether the foldable IGORF was the first to be selected and whether selection has only retained the combinations of IGORFs that do not affect the foldability of the preexisting selected product are exciting questions that deserve further investigations.

**Figure 4.**
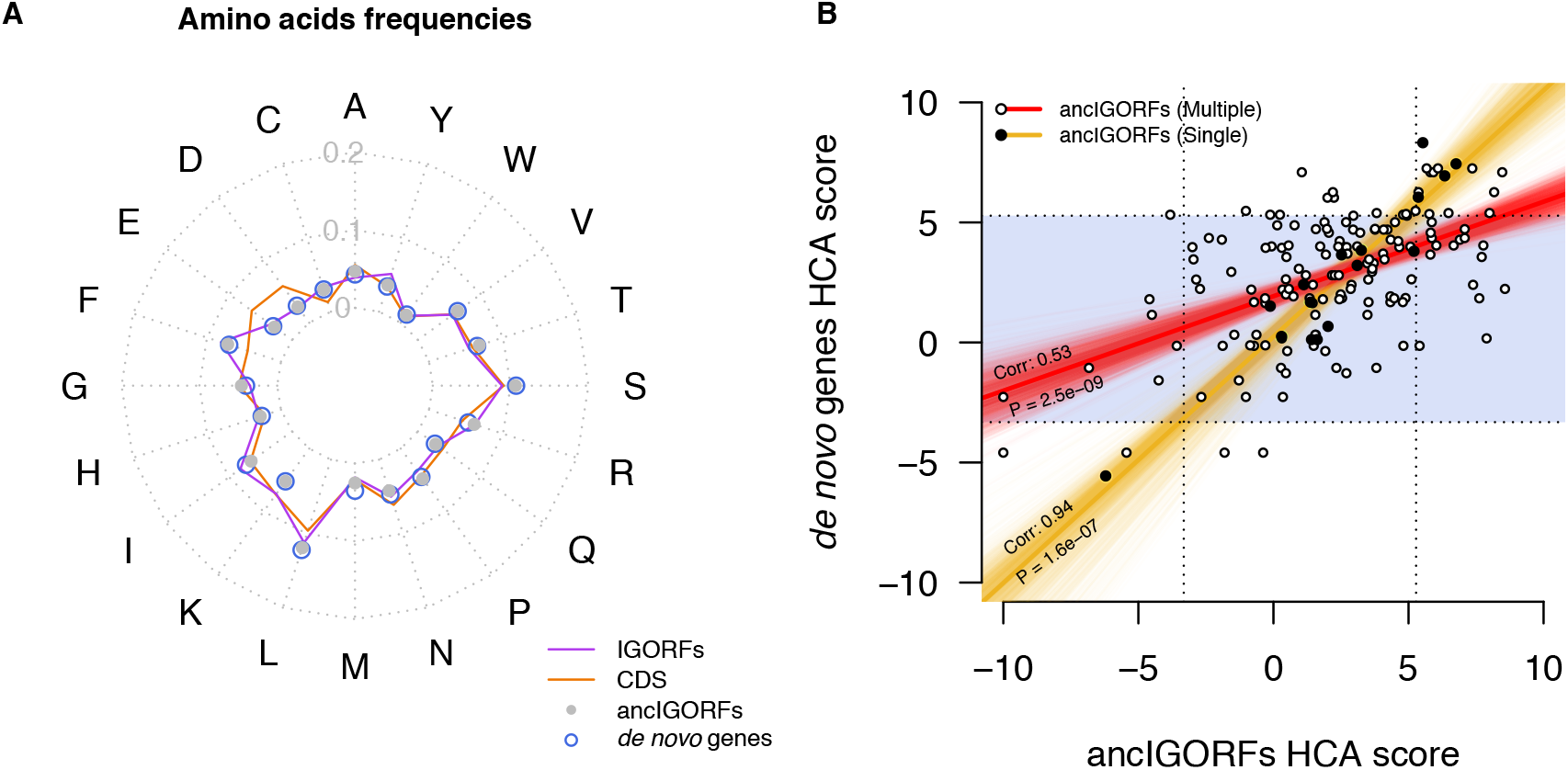
(A) Radar plot reflecting the 20 amino acid frequencies of IGORFs, ancIGORFs, *de novo* genes and CDS. (B) The fold potential of a single ancIGORF *de novo* gene is mostly determined by the one of its parent ancIGORFs while the combination of several ancIGORFs through frameshift events and STOP codon mutations lead most of the time to a foldable product. Plot of the HCA score of each *de novo* gene with those of its parent ancIGORF(s). Single ancIGORF and multiple ancIGORFs *de novo* genes are represented by black and white points respectively. The linear regression of single and multiple ancIGORFs de novo genes’ HCA scores versus the score of their parent ancIGORF(s) are represented in yellow and red respectively. The Pearson correlation coefficients and the corresponding p-values are indicated on the plot. The blue region indicates *de novo* genes encoding proteins predicted as foldable.

### Translated and ancestral IGORFs display intermediate properties between IGORFs and CDS

Next, we performed two ribosome profiling experiments on *S. cerevisiae* (strain BY4742) and used additional ribosome profiling data from three other experiments to define two types of translated IGORFs (Radhakrishnan et al. 2016; Thiaville et al. 2016). The former corresponds to IGORFs that are occasionally translated with a weak translation signal (at least 10 reads in one experiment - see Methods for more details). They are mostly expected to be short-lived in evolutionary history, though some of them may give rise to future novel genes. The latter corresponds to IGORFs with a strong translation signal (more than 30 reads in at least two experiments) and whose translation is strongly favored over the overlapping IGORFs in the other phases (i.e. selectively translated IGORFs)(see Methods for more details). This suggests the optimization of their translation activity which could be related to the emergence of a functional translation product. We identified 1235 occasionally translated IGORFs and 31 selectively translated IGORFs. Figure 5 shows the boxplot distributions of the sizes of the sequences, clusters and linkers of the translated IGORFs and all other ORF categories (e.g. IGORFs, ancIGORFs, *de novo* genes and CDS) along with their number of clusters per sequence. In line with Carvunis et al. (2012), the Figure 5 reveals for most properties, a continuum from IGORFs to CDS reflecting the successive stages preceding the emergence of a *de novo* gene until the establishment of a genuine gene. Interestingly, the selectively translated IGORFs and the ancIGORFs are both longer than IGORFs (Mann-Whitney U-test, P =3.4e-02 and 1.3e-22 respectively) and display longer linkers (Mann-Whitney U-test, P =2.6e-02 and 1.8e-02), though the effect is less pronounced for the 31 translated IGORFs (Fig. 5). This can be explained by the fact that among the latter, only a handful of them will give rise to a *de novo* gene whereas all ancIGORF have indeed given rise to a *de novo* gene.

**Figure 5.**
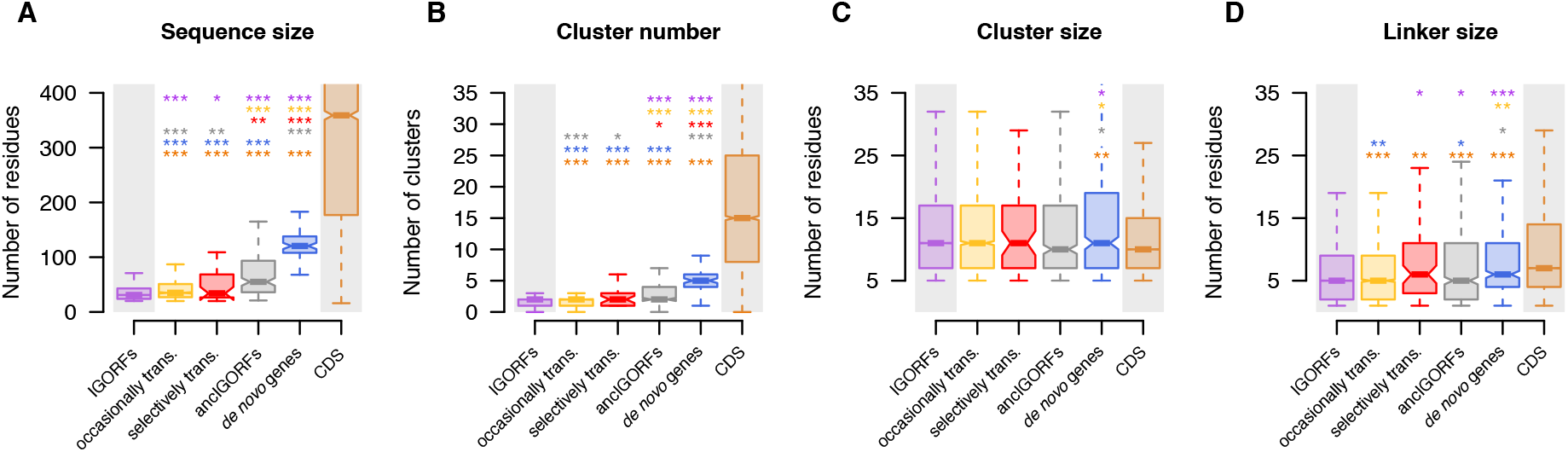
Continuum of sequence and structural properties between the early stages of *de novo* gene emergence and CDS. Comparison of (A) the sequence size, (B) cluster number, (C) cluster sizes, and (D) linker sizes for each ORF categories (IGORFs in purple, Occasionally translated IGORFs in yellow, Selectively translated IGORFs in red, ancIGORFs in grey, *de novo* genes in blue and CDS in orange). The p-values were computed with the Mann-Whitney U-test (one-sided for (A), (B), (D) and two-sided for (C)). Asterisks denote level of significance: *p < 0.05, **p < 0.01, ***p < 0.001. For each boxplot, the color of the asterisks indicates the ORF category used for the comparison. The exact p-values are given in tables S3-6.

Strikingly, the HCA cluster size remains invariant for all ORF categories except the one of *de novo* gene clusters, thereby reinforcing the concept of hydrophobic clusters as elementary building blocks of proteins (Fig. 5). The increase in *de novo* gene cluster size cannot be explained by the hydrophobic content of *de novo* genes which is similar to those of IGORFs and ancIGORFs (Fig. 4A). However, we hypothesize that longer clusters mostly result from the fusion of IGORFs through STOP codon mutations or frameshift events as observed in the example of the YPR126C *de novo* gene (Figure S8A).

Interestingly, this gene results from the fusion of three ancIGORFs through STOP codon mutations which led to longer clusters in YPR126C. Similarly, the fusion of ancIGORFs can also give rise to longer linkers as observed with the YMR153C-A *de novo* gene (Fig. S8B). The fact that CDS are characterized by longer linkers while their cluster size is similar to the one of IGORFs suggests that harboring long linkers is a criterion that has been selected over evolution whereas it is not the case for long clusters. Having shown that CDS are enriched in hydrophilic residues (Fig S2), we can hypothesize that the mutations of hydrophobic residues toward hydrophilic ones can disrupt long clusters or can switch cluster extremities into linker extremities, thereby decreasing their size over time.

## Discussion

In this work, we showed that the noncoding genome encodes the raw material for making proteins. In particular, we showed the widespread existence in the noncoding genome of the elementary building blocks of protein structures which consist of hydrophobic clusters that have been shown to be associated in protein coding sequences with regular secondary structures (Bitard-Feildel and Callebaut 2017; Bitard-Feildel et al. 2018; Lamiable et al. 2019). We showed that hydrophobic clusters in noncoding sequences display sizes similar to those observed in CDS and that ancestral IGORFs that gave birth to *de novo* genes are characterized by a larger number of clusters (Fig. 5). In contrast, CDS are enriched in longer linkers which probably contribute to optimize the local arrangements of secondary structures, provide flexibility to proteins, and specificity in protein interactions. This observation is in line with several studies reporting a central role to loops in protein function and structural innovation (Blouin et al. 2004; Espadaler et al. 2006; Tendulkar et al. 2004; Papaleo et al. 2016). Like Schmitz et al. (2018), we stipulate that the increase in intrinsic structural disorder observed for old genes by Carvunis et al. (2012), is related to the fact that CDS are characterized by longer linkers, thereby inducing inevitably an increase in the disorder score. As a matter of fact, most CDS display HCA scores similar to those of globular proteins, with low disorder propensities (Fig. 2). Overall, we showed an enrichment in polar and charged residues for both linkers and clusters of CDS which is likely accompanied by an increase in specificity of protein folds and interactions through the optimization of the folding and assembly processes (Lumb and Kim 1995).

Nevertheless, how a noncoding sequence becomes coding remains unclear. In this work, we propose the IGORFs as elementary modules of protein birth and evolution. IGORFs can serve as starting points for *de novo* gene emergence or can be combined together, thus increasing protein sizes, contributing to protein modularity, and leading to more complex protein architectures. They recall the short protein fragments, reported so far, that result from different protein structure decompositions (Nepomnyachiy et al. 2017; Papandreou et al. 2004; Alva et al. 2015; Postic et al. 2017; Kolodny et al. 2020; Berezovsky et al. 2000, 2001; Lamarine et al. 2001). Interestingly, we showed that these elementary modules encompass all the protein structural diversity observed in CDS. A majority of IGORFs encode peptides predicted as foldable while an important fraction displays high HCA scores and aggregation propensities. Some of the latter, though not the majority (28%), are predicted with at least one TM domain and may “safely” locate in membranes as proposed in Vakirlis et al. (2020). The impact of the other high HCA score IGORFs on the cell deserves further investigations. Nevertheless, we can hypothesize that if produced, most of the time, their concentration will not be sufficient to be deleterious (Langenberg et al. 2020). Indeed, it seems that for CDS, a certain degree of aggregation is tolerated at low concentration (see Fig. S9). On the other hand, although IGORFs with intermediate HCA scores may exhibit a certain propensity for aggregation, we can hypothesize that these aggregation-prone regions, under the hydrophobic effect, may play a role in their capacity to fold, in line with the hypothesis of an amyloid origin of the globular proteins (Langenberg et al. 2020; Greenwald and Riek 2012). Indeed, the balanced equilibrium of hydrophobic and hydrophilic residues observed for these IGORFs (39.1% of hydrophobic residues to be compared with the 50.8% observed for high HCA score IGORFs) may render possible the burying of aggregation-prone regions and the exposure of hydrophilic residues that is accompanied by an increase in foldability. We can hypothesize that, if produced, these IGORFs could form small compact structures and/or could be stabilized through oligomerization or interactions with other proteins. Precisely, we showed that ancestral IGORFs predating *de novo* gene emergence are not IGORF-like, but rather enriched in sequences with a high propensity for foldability. This reveals that at least for *S. cerevisiae*, foldable IGORFs are more likely to give rise to novel genes, though, it must be confirmed for other lineages with different GC contents (Foy et al. 2019; Basile et al. 2017; Heames et al. 2020). Nevertheless, we can reasonably hypothesize that *de novo* peptides struggle to fold to a well-defined and specific 3D structure as shown with the young *de novo* gene BSC4 identified in the *S. cerevisiae* lineage (Bungard et al. 2017; Namy et al. 2003). Recently, Bungard et al. (2017) reported that the Bsc4 protein folds partially to an ordered structure that is unlikely to be unfolded according to Circular Dichroism spectra and bioinformatic analyses. Interestingly, despite this “rudimentary” fold, they show through Mass Spectrometry and denaturation experiments that Bsc4 is able to form compact oligomers. Consistently with Bungard et al. (2017), HCA predicts Bsc4 as foldable with an intermediate HCA score of 1.98 (38% of hydrophobic residues), though it cannot predict whether Bsc4 folds completely or partially to an ordered structure. Overall, except the sequence length, the sequence and structural features of Bsc4 are similar to those of ancIGORFs. This suggests that similarly to the Bsc4 protein, young *de novo* proteins can optimize their fold specificity as well as their interactions with their environment through amino acid substitutions toward hydrophilic residues, thereby leading to the amino acid composition and the well-defined structures of most canonical proteins.

Altogether, these results enable us to propose a model (Fig. 6) which gives a central role to IGORFs in *de novo* gene emergence and to a lesser extent in protein evolution, thus completing the large palette of protein evolution mechanisms such as duplication events, horizontal gene transfer, domain shuffling... Once an IGORF is selected, it can elongate through frameshift events and/or STOP codon mutations, thus incorporating a neighboring IGORF (Fig. 6B). Bartonek et al. (2020), showed that the hydrophobicity profiles of protein sequences remain invariant after frameshift events. Consequently, frameshift events are most of the time expected to incorporate an IGORF that encodes a peptide with a hydrophobicity profile similar to that of the preexisting gene. This suggests that the fold potential is a critical feature that needs to be conserved even in noncoding sequences, being preserved in +1, −1 phases through the structure of the genetic code. In addition, we showed that combining IGORFs encoding for foldable peptides with IGORFs encoding for disorder or aggregation-prone peptides has low impact on the foldability of the resulting *de novo* protein (Fig. 4B). We can hypothesize that the newly integrated IGORFs will benefit from the structural properties of the preexisting IGORF network. More generally, proteins can be seen as assemblies on an ancient protein core, whatever its evolutionary history, of either duplicated, shuffled domains or *de novo* translated products encoded by neighboring IGORFs (Fig. 6D). Our model is supported by previous observations which show that (i) *de novo* genes are shorter than old ones (Tautz and Domazet-Lošo 2011; Wolf et al. 2009), (ii) the size of *de novo* gene exons are similar to those of old genes (Schlötterer 2015; Neme and Tautz 2013; Palmieri et al. 2014), and (iii) novel domains are generally observed in the C-terminal regions (Bornberg-Bauer et al. 2015; Klasberg et al. 2018). Nevertheless, the increase in linker sizes observed between the different ORF categories remains unclear. It is unknown whether harboring long linkers is accompanied by an increase in foldability and is thus a selected criterion as suggested by the observation that ancIGORFs and selectively translated IGORFs display longer linkers than IGORFs in general. Also, mutational events such as amino acid mutations toward hydrophilic residues and frameshift events or STOP codon mutations may result in longer linkers (Fig. 6BC) as observed in the example of the YMR153C-A *de novo* gene (Fig. S8B).

**Figure 6.**
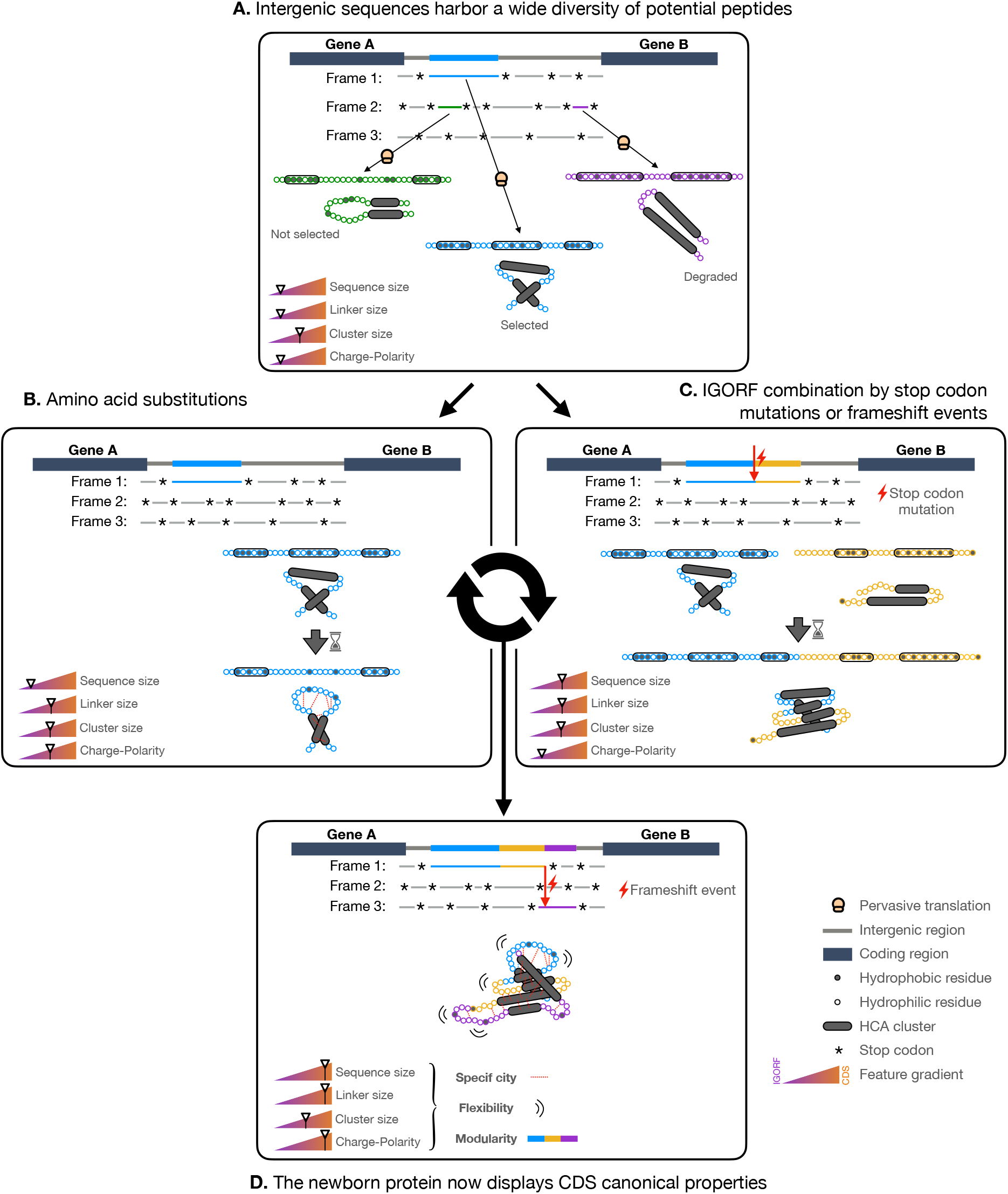
Model of *de novo* gene emergence and protein evolution with IGORFs as elementary structural modules. (A) IGORFs encode a wide diversity of peptides from disorder prone to aggregation prone ones, among which, a vast amount is expected to be able to fold in solution. Upon pervasive translation, some peptides that can be deleterious or not, will be degraded right away. Among the others, the blue one will confer an advantage to the organism and will be further selected, thus providing a starting point for *de novo* gene birth. (B) The STOP codon of the starting point IGORF can be mutated into an amino acid, thereby adding the yellow IGORF to the preexisting selected IGORF and elongating its size. The fusion of the two IGORFs leads to the creation of a longer linker at the IGORFs’ junction. This can also apply for hydrophobic clusters which can elongate through frameshift events and STOP codon mutations, though it seems that the resulting long clusters observed in *de novo* genes are not selected in CDS. Amino acid mutations toward hydrophilic residues may disrupt long clusters into smaller ones or redefine their borders by converting their extremities into hydrophilic residues (C) The starting point IGORF, once selected, is subjected to amino acid substitutions thereby increasing the overall proportion of hydrophilic residues of the encoded peptide. In the present case, this induces (i) the disruption of the second cluster resulting to the increase of the size of the central linker and (ii) the establishment of specific interactions between hydrophilic residue (red dots) which increase the specificity of the folding process and the resulting fold (D) After multiple events of amino acid substitutions and IGORF combinations through STOP codon mutations or frameshift events, we obtain a protein which displays the canonical features of CDS (i.e. long sequences, long linkers, enrichment in polar and charged residues) which enable the optimization of its flexibility, the increase in specificity of its folding process, 3D fold and interactions and finally participate along with domain shuffling or duplication events in the modular architecture of genuine proteins.

In this work, we propose a model that covers the genesis of all the structural diversity observed in current proteins. Although, we showed that IGORFs encoding foldable peptides are more likely to give rise to novel genes, disordered or aggregation-prone *de novo* proteins may emerge occasionally (see squares bottom-left and top-right in Fig. 4B). Particularly, in line with Vakirlis et al. (2020), we observe an enrichment in TM-prone sequences for ancIGORFs compared to IGORFs (26.3% and 16.1% respectively, one proportion z test, P = 2e-4), although the majority of peptides encoded by ancIGORFs are not TM-prone. More interestingly, we observe a strong correlation between the foldability propensity of the single ancIGORFs and their resulting *de novo* proteins (Fig. 4B) and that disorder or aggregation prone *de novo* proteins are most of the time (79%) associated with ancIGORFs expected to encode disordered or aggregation-prone peptides as well, suggesting that the structural properties of *de novo* proteins are already encoded in the ancestral peptide they originate from (see in Fig. 3, the blue dots connected by green and pink lines in the low and high HCA score categories respectively). Whether the fold potential of a starting point IGORF conditions the structural properties of the resulting *de novo* protein is an exciting question that deserves further studies. Indeed, we can hypothesize that once an IGORF is selected, it can elongate over time through the incorporation of neighboring IGORFs, provided that the latter do not affect the fold potential of the preexisting protein. In accordance with Vakirlis et al. (2020), we can reason that once a starting point IGORF is selected, it engenders novel selected effects which in turn, increase the constraints exerted on it and subsequently reduce the possibility of future changes. It is thus tempting to speculate that the structural properties of the peptide encoded by the starting point IGORF will be retained during evolution through the elimination of the deleterious IGORFs’ combinations. All these observations suggest that the fold diversity observed in current proteins has been originally inherited from the diversity of the fold potential already encoded in the noncoding genome.

## Methods

### Datasets

#### CDS and IGORFs

The CDS were extracted from the genome of *Saccharomyces cerevisiae* S288C according to the genome annotation of the S*accharomyces* Genome database (Cherry et al. 2012). All unannotated ORFs of at least 60 nucleotides, no matter if they start with an AUG codon, were extracted from the 16 yeast chromosomes. 60 nucleotides correspond to 20 amino acids which is a reasonable minimum size for a peptide to acquire its own fold (Qiu et al. 2002). The hydrophobicity profiles of overlapping sequences in two different frames were shown to be correlated in Bartonek et al. (2020). Therefore, in order to prevent any bias from CDS hydrophobicity profiles, we only retained ORFs that are free from overlap with another gene or that partially overlap with a gene if the non-overlapping region is more than 70% of the IGORFs sequence.

#### Datasets of reference

The disorder dataset consists of 731 disordered regions extracted from intrinsically disordered proteins of the Disprot database (Hatos et al. 2020), that were used for the calibration of HCAtk (Bitard-Feildel and Callebaut 2018). The globular dataset consists of 559 globular proteins extracted from the Protein Data Bank (Berman et al. 2000; Burley et al. 2021) that were used for the calibration of IUPred (Dosztanyi et al. 2005; Dosztányi 2018; Mészáros et al. 2018, 2009). The transmembrane regions dataset gathers 1269 transmembrane regions extracted from the transmembrane proteins contained in the PDBTM database (Tusnády et al. 2004, 2005; Kozma et al. 2012), thereby expected to form aggregates in solution. We only retained transmembrane segments longer than 20 amino acids corresponding to the minimum size of an IGORF. These TM regions only match buried regions of TM proteins and are not expected to display the same sequence and structural properties as the complete membrane proteins they were extracted from. Indeed, membrane proteins including integral membrane proteins which involve TM domains along with extracellular or cytosolic domains of variable sizes.

### Estimation of the fold potential, the aggregation, disorder and TM propensities

The foldability potential was estimated using a score derived from the HCA (Hydrophobic Cluster Analysis) approach using the HCAtk (Bitard-Feildel et al. 2018; Bitard-Feildel and Callebaut 2018). HCA divides a protein sequence into (i) clusters gathering strong hydrophobic residues (V, I, L, F, M, Y, W) or cysteines, and (ii) linkers composed of at least 4 non-hydrophobic residues (or a proline). As supported by analysis of experimental 3D structures, hydrophobic clusters match regular secondary structures (single ones or more, if separated by short loops) while the linkers indicate flexible regions generally corresponding to loops. The fold potential of a sequence is determined by its density in hydrophobic clusters but also by the density of hydrophobic amino acids within hydrophobic clusters. It is reflected with the HCA score which ranges from −10 to +10. Low scores indicate sequences depleted in hydrophobic clusters, which are likely to be disordered whereas high scores are associated with a very high density in hydrophobic clusters, that are expected to form aggregates in solution. The aggregation propensity of a sequence was assessed with TANGO (Fernandez-Escamilla et al. 2004; Linding et al. 2004; Rousseau et al. 2006a). Following the criteria presented in Linding et al. (2004), a residue was considered as participating in an aggregation prone region if it was located in a segment of at least five consecutive residues which were predicted as populating a b-aggregated conformation for more than 5%. Then, the aggregation propensity of each sequence is defined as the fraction of residues predicted in aggregation prone segments. The disorder propensity was probed with IUPred (Dosztanyi et al. 2005; Dosztányi 2018; Mészáros et al. 2018, 2009) using the short prediction option. To be consistent with the criteria used for assessing the aggregation propensity, we considered a residue as participating in a disordered region if it is located in a segment of at least five consecutive residues, each presenting a disorder probability higher than 0.5. Then, the disorder propensity of each sequence is defined as the fraction of residues predicted in disordered prone segments. The presence of at least one TM domain was predicted with TMHMM (Krogh et al. 2001).

### Protein abundances and amino acid propensities

Protein abundance data were extracted from the PaxDB database (Wang et al. 2012). In order to depict the impact of the avoidance of nonspecific interactions with the ribosome, we only retained cytoplasmic proteins as annotated in Uniprot^35^. The propensity of an amino acid *i* to be found in a CDS is defined by the log ratio of the frequencies of the amino acid *i* in CDS versus IGORFs as follows:

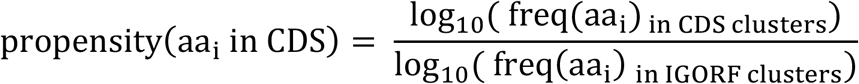

### Reconstruction of Ancestral IGORFs

To reconstruct the ancIGORFs of *S. cerevisiae*, we used the genomes of the neighboring species *S. paradoxs* (Durand et al. 2019), *S. arboricola* (Yue et al. 2017), *S. mikatae, S. kudriavzevii,* and *S. uvarum* (Scannell et al. 2011). Based on four independent studies which each listed *de novo* genes of the *S. cerevisiae* S288C genome, we retained all *de novo* genes identified in at least two studies (Carvunis et al. 2012; Vakirlis et al. 2018; Lu et al. 2017; Wu and Knudson 2018). This led to a total of 171 *de novo* genes among which we retained those (see the list of the 70 *de novo* genes in Table S2) for which we were able to identify at least two additional homologous sequences in the neighboring species among which at least one had to be noncoding in order to reconstruct the corresponding nongenic region in the ancestor. Therefore, we searched for the orthologous genes of the 70 *de novo* genes in the neighboring species using Blast (evalue < 1e-2) (Fig. S5A). Then, based on the species tree presented in Figure S5A and starting from the branch of *S. cerevisiae*, we traced back to the root and identified the first node branching with a branch for which no orthologous gene had been detected (yellow circle in Fig. S5A). We can reasonably hypothesize that the corresponding locus in the ancestor was still nongenic. We then searched for the corresponding nongenic regions in the remaining species with tblastn (evalue < 1e-2). Then following the protocol described by Vakirlis and McLysaght (2019), the resulting homologous nucleotide sequences and orthologous *de novo* genes were subsequently aligned with MACSE v2.05 (Ranwez et al. 2011, 2018) and the corresponding phylogenetic tree was constructed with PHYML (Guindon et al. 2010). The multiple sequence alignment and its corresponding tree were given as input to PRANK (Löytynoja and Goldman 2010) for the reconstruction of the corresponding ancestral nongenic nucleotide sequence (Fig. S5BC). Finally, the ancestral nucleotide sequences were translated into the three reading frames. The resulting IGORFs were then aligned with the *de novo* gene of *S. cerevisiae* with LALIGN (Huang and Miller 1991) and those sharing a homology with it were extracted, the others were eliminated (Fig. S5D).

### Ribosome Profiling analyses

#### Ribosome profiling experiments

Cells were grown overnight in 0.5 liter of liquid glucose-YPD till an OD_600_ of 0.6, 50 microg/microl of cycloheximide were added to the culture and incubated during 5 min and kept at + 4°C. The pellet of yeast cells was recovered by centrifugation during 5 min at 5000 rpm in Beckman F10 rotor at + 4°C. Total RNA and polysomes were extracted as previously described (Baudin-Baillieu et al. 2014). Briefly, cells were lysated by vortex during 15 min in 500 microl of polysome buffer (10 mM Tris-acetate pH7.5; 0.1M NaCl and 30 mM Mg-acetate) in presence of glass beads in Eppendorf tube, followed by 5 min of centrifugation at 16 krcf at + 4°C. Ribosome-protected mRNA fragments (RPFs) were generated by the treatment following the ratio of 1 OD260nm of extract with 15 U of RNase I during 1 h at 25°C. Monosomes were collected by 2h15 min centrifugation on a 24% sucrose cushion at + 4°C on TLA 110 rotor at 110 krpm. The monosomes were resuspended with 500 microl of polysome buffer. RNA was purified by phenol–chloroform extraction and 28-34 nucleotides RPFs were recovered by electrophoresis in a 17% acrylamide (19/1) 7M urea in 1 × TAE gel. These RPFs were depleted of ribosomal RNA by treatment with the Ribo-Zero Gold rRNA removal kit for yeast from Illumina company. RPF libraries were generated with NEBNext Small RNA Sample Prep Kit, according to the manufacturer’s protocol, and were checked with the bioanalyser small RNA kit. Sequencing was performed by a HighSeq 2000 (Illumina) 75-nucleotide single-read protocol.

#### Additional ribosome profiling data

we used three additional experiments that were retrieved by Radhakrishnan et al. (2016) (GEO accession numbers GSM2147982 and GSM2147983) and Thiaville et al. (2016) (GEO accession number GSM1850252).

#### Selection of RPF (Ribosome Protected Fragments)

Ribosome profiling reads were mapped on the genome of *S. cerevisiae* S288C using Bowtie (Langmead et al. 2009). For this study we kept only the 28-mers since on average 90% of them were mapped on a CDS in the correct reading frame (i.e. are in frame with the start codon of the CDS) (see Fig. S10-left).

#### Periodicity

The periodicity is calculated using a metagene profile. It provides the number of footprints relative to all annotated start codons in a selected window. The metagene profile is obtained by pooling together all the annotated CDS and counting the number of RPFs at each nucleotide position (determined by the site P of each 28-mer). Results presented in Figure S10 show a clear accumulation of signal over the CDS, and a nice periodicity over the 100 first nucleotides.

#### Identification of the occasionally translated IGORFs

we retained the IGORFs with at least 10 reads in at least one experiment.

#### Identification of the selectively translated IGORFs

we kept the IGORFs with at least 30 reads in at least two experiments and for which the fraction of reads in the frame of the IGORF was higher than 0.8, reflecting that the translation of the IGORF is favored over the other overlapping ones.

### Statistical analyses

All statistical analyses that aimed at comparing distributions were performed in R using the Kolmogorov–Smirnov test (two-sided) when comparing whether the HCA score distributions are statistically different and the Mann Whitney U-test for the comparison of the median cluster size, linker size, sequence size and cluster number distributions (bilateral test for the comparison of cluster sizes and unilateral test for the other properties). We used the one proportion z test for the comparison of the proportion of disordered, foldable or aggregation prone sequences between different ORF categories. In order to (i) circumvent the *p-value problem* inherent to large samples (i.e. extremely large samples such as the one of IGORFs induce low p-values) (Lin et al. 2013), tests were performed iteratively 1000 times on samples of 500 individuals randomly chosen from the initial sample when it was larger than 500 individuals. The averaged p-value over the 1000 iterations was subsequently calculated.

## Availability of data

The raw ribosome profiling data are currently being deposited on the NCBI GEO platform but are available for referees at: http://bim.i2bc.paris-saclay.fr/anne-lopes/data_Papadopoulos/Riboseq_data_Papadopoulos/ Raw and calculated data along with custom scripts used in this study are available as Supplemental Data files. The extraction of IGORFs and their structural properties (foldability potential, disorder and aggregation propensities) were calculated using our in-house programs (ORFtrack and ORFold respectively) available in the ORFmine package at: https://github.com/i2bc/ORFmine

## Supplemental Material

Supplemental Figures are stored in Papadopoulos_Supplemental_Figures.pdf

Supplemental Tables are stored in Papadopoulos_Supplemental_Tables.pdf

## Funding

CP work was supported by a French government fellowship.

## Competing Interest Statement

The authors declare no conflict of interest.

## Author contributions

CP, MR, IH performed research, CP, MR, IH, ON, AL analyzed data. CP, AL designed research. CP, IC, JCG, ON, OL, AL wrote the paper. AL conceived the project.

## Notes

### Competing Interest Statement

The authors have declared no competing interest.

## References

Alva V, Söding J, Lupas AN. 2015. A vocabulary of ancient peptides at the origin of folded proteins. Elife 4: e09410.

Bartonek L, Braun D, Zagrovic B. 2020. Frameshifting preserves key physicochemical properties of proteins. Proc Natl Acad Sci 117: 5907–5912.

Basile W, Sachenkova O, Light S, Elofsson A. 2017. High GC content causes orphan proteins to be intrinsically disordered. PLoS Comput Biol 13: e1005375.

Baudin-Baillieu A, Legendre R, Kuchly C, Hatin I, Demais S, Mestdagh C, Gautheret D, Namy O. 2014. Genome-wide translational changes induced by the prion [PSI+]. Cell Rep 8: 439–448.

Berezovsky IN, Grosberg AY, Trifonov EN. 2000. Closed loops of nearly standard size: common basic element of protein structure. Febs Lett 466: 283–286.

Berezovsky IN, Kirzhner VM, Kirzhner A, Trifonov EN. 2001. Protein folding: looping from hydrophobic nuclei. Proteins Struct Funct Bioinforma 45: 346–350.

Berman HM, Westbrook J, Feng Z, Gilliland G, Bhat TN, Weissig H, Shindyalov IN, Bourne PE. 2000. The protein data bank. Nucleic Acids Res 28: 235–242.

Bitard-Feildel T, Callebaut I. 2017. Exploring the dark foldable proteome by considering hydrophobic amino acids topology. Sci Rep 7: 1–13.

Bitard-Feildel T, Callebaut I. 2018. HCAtk and pyHCA: A Toolkit and Python API for the Hydrophobic Cluster Analysis of Protein Sequences. bioRxiv 249995.

Bitard-Feildel T, Heberlein M, Bornberg-Bauer E, Callebaut I. 2015. Detection of orphan domains in Drosophila using “hydrophobic cluster analysis.” Biochimie 119: 244–253.

Bitard-Feildel T, Lamiable A, Mornon J, Callebaut I. 2018. Order in disorder as observed by the “hydrophobic cluster analysis” of protein sequences. Proteomics 18: 1800054.

Blevins WR, Ruiz-Orera J, Messeguer X, Blasco-Moreno B, Villanueva-Cañas JL, Espinar L, Díez J, Carey LB, Albà MM. 2021. Uncovering de novo gene birth in yeast using deep transcriptomics. Nat Commun 12: 1–13.

Blouin C, Butt D, Roger AJ. 2004. Rapid evolution in conformational space: a study of loop regions in a ubiquitous GTP binding domain. Protein Sci 13: 608–616.

Bornberg-Bauer E, Alba MM. 2013. Dynamics and adaptive benefits of modular protein evolution. Curr Opin Struct Biol 23: 459–466.

Bornberg-Bauer E, Schmitz J, Heberlein M. 2015. Emergence of de novo proteins from ‘dark genomic matter’by ‘grow slow and moult.’ Biochem Soc Trans 43: 867–873.

Bungard D, Copple JS, Yan J, Chhun JJ, Kumirov VK, Foy SG, Masel J, Wysocki VH, Cordes MH. 2017. Foldability of a natural de novo evolved protein. Structure 25: 1687–1696.

Burley SK, Bhikadiya C, Bi C, Bittrich S, Chen L, Crichlow GV, Christie CH, Dalenberg K, Di Costanzo L, Duarte JM. 2021. RCSB Protein Data Bank: powerful new tools for exploring 3D structures of biological macromolecules for basic and applied research and education in fundamental biology, biomedicine, biotechnology, bioengineering and energy sciences. Nucleic Acids Res 49: D437–D451.

Carvunis A-R, Rolland T, Wapinski I, Calderwood MA, Yildirim MA, Simonis N, Charloteaux B, Hidalgo CA, Barbette J, Santhanam B. 2012. Proto-genes and de novo gene birth. Nature 487: 370–374.

Cherry JM, Hong EL, Amundsen C, Balakrishnan R, Binkley G, Chan ET, Christie KR, Costanzo MC, Dwight SS, Engel SR. 2012. Saccharomyces Genome Database: the genomics resource of budding yeast. Nucleic Acids Res 40: D700–D705.

Dosztányi Z. 2018. Prediction of protein disorder based on IUPred. Protein Sci 27: 331–340.

Dosztanyi Z, Csizmok V, Tompa P, Simon I. 2005. The pairwise energy content estimated from amino acid composition discriminates between folded and intrinsically unstructured proteins. J Mol Biol 347: 827–839.

Durand É, Gagnon-Arsenault I, Hallin J, Hatin I, Dubé AK, Nielly-Thibault L, Namy O, Landry CR. 2019. Turnover of ribosome-associated transcripts from de novo ORFs produces gene-like characteristics available for de novo gene emergence in wild yeast populations. Genome Res 29: 932–943.

Ekman D, Elofsson A. 2010. Identifying and quantifying orphan protein sequences in fungi. J Mol Biol 396: 396–405.

Espadaler J, Querol E, Aviles FX, Oliva B. 2006. Identification of function-associated loop motifs and application to protein function prediction. Bioinformatics 22: 2237–2243.

Faure G, Callebaut I. 2013. Comprehensive repertoire of foldable regions within whole genomes. PLoS Comput Biol 9.

Fernandez-Escamilla A-M, Rousseau F, Schymkowitz J, Serrano L. 2004. Prediction of sequence-dependent and mutational effects on the aggregation of peptides and proteins. Nat Biotechnol 22: 1302–1306.

Foy SG, Wilson BA, Bertram J, Cordes MH, Masel J. 2019. A shift in aggregation avoidance strategy marks a long-term direction to protein evolution. Genetics 211: 1345–1355.

Ganesan A, Siekierska A, Beerten J, Brams M, Van Durme J, De Baets G, Van der Kant R, Gallardo R, Ramakers M, Langenberg T. 2016. Structural hot spots for the solubility of globular proteins. Nat Commun 7: 1–15.

Greenwald J, Riek R. 2012. On the possible amyloid origin of protein folds. J Mol Biol 421: 417–426.

Guindon S, Dufayard J-F, Lefort V, Anisimova M, Hordijk W, Gascuel O. 2010. New algorithms and methods to estimate maximum-likelihood phylogenies: assessing the performance of PhyML 3.0. Syst Biol 59: 307–321.

Hatos A, Hajdu-Soltész B, Monzon AM, Palopoli N, Álvarez L, Aykac-Fas B, Bassot C, Benítez GI, Bevilacqua M, Chasapi A. 2020. DisProt: intrinsic protein disorder annotation in 2020. Nucleic Acids Res 48: D269–D276.

Heames B, Schmitz J, Bornberg-Bauer E. 2020. A continuum of evolving de novo genes drives protein-coding novelty in Drosophila. J Mol Evol 88: 382–398.

Huang X, Miller W. 1991. A time-efficient, linear-space local similarity algorithm. Adv Appl Math 12: 337–357.

Ingolia NT, Lareau LF, Weissman JS. 2011. Ribosome profiling of mouse embryonic stem cells reveals the complexity and dynamics of mammalian proteomes. Cell 147: 789–802.

Klasberg S, Bitard-Feildel T, Callebaut I, Bornberg-Bauer E. 2018. Origins and structural properties of novel and de novo protein domains during insect evolution. FEBS J 285: 2605–2625.

Kolodny R, Nepomnyachiy S, Tawfik DS, Ben-Tal N. 2020. Bridging themes: short protein segments found in different architectures. bioRxiv.

Kozma D, Simon I, Tusnady GE. 2012. PDBTM: Protein Data Bank of transmembrane proteins after 8 years. Nucleic Acids Res 41: D524–D529.

Krogh A, Larsson B, Von Heijne G, Sonnhammer EL. 2001. Predicting transmembrane protein topology with a hidden Markov model: application to complete genomes. J Mol Biol 305: 567–580.

Lamarine M, Mornon J-P, Berezovsky IN, Chomilier J. 2001. Distribution of tightened end fragments of globular proteins statistically matches that of topohydrophobic positions: towards an efficient punctuation of protein folding? Cell Mol Life Sci CMLS 58: 492–498.

Lamiable A, Bitard-Feildel T, Rebehmed J, Quintus F, Schoentgen F, Mornon J-P, Callebaut 2019. A topology-based investigation of protein interaction sites using Hydrophobic Cluster Analysis. Biochimie 167: 68–80.

Langenberg T, Gallardo R, van der Kant R, Louros N, Michiels E, Duran-Romaña R, Houben B, Cassio R, Wilkinson H, Garcia T. 2020. Thermodynamic and evolutionary coupling between the native and amyloid state of globular proteins. Cell Rep 31: 107512.

Langmead B, Trapnell C, Pop M, Salzberg SL. 2009. Ultrafast and memory-efficient alignment of short DNA sequences to the human genome. Genome Biol 10: 1–10.

Levy ED, De S, Teichmann SA. 2012. Cellular crowding imposes global constraints on the chemistry and evolution of proteomes. Proc Natl Acad Sci 109: 20461–20466.

Lin M, Lucas HC Jr, Shmueli G. 2013. Research commentary–too big to fail: large samples and the p-value problem. Inf Syst Res 24: 906–917.

Linding R, Schymkowitz J, Rousseau F, Diella F, Serrano L. 2004. A comparative study of the relationship between protein structure and β-aggregation in globular and intrinsically disordered proteins. J Mol Biol 342: 345–353.

Löytynoja A, Goldman N. 2010. webPRANK: a phylogeny-aware multiple sequence aligner with interactive alignment browser. BMC Bioinformatics 11: 579.

Lu T-C, Leu J-Y, Lin W-C. 2017. A comprehensive analysis of transcript-supported de novo genes in Saccharomyces sensu stricto yeasts. Mol Biol Evol 34: 2823–2838.

Lumb KJ, Kim PS. 1995. A buried polar interaction imparts structural uniqueness in a designed heterodimeric coiled coil. Biochemistry 34: 8642–8648.

Macossay-Castillo M, Marvelli G, Guharoy M, Jain A, Kihara D, Tompa P, Wodak SJ. 2019. The balancing act of intrinsically disordered proteins: enabling functional diversity while minimizing promiscuity. J Mol Biol 431: 1650–1670.

Mészáros B, Erdős G, Dosztányi Z. 2018. IUPred2A: context-dependent prediction of protein disorder as a function of redox state and protein binding. Nucleic Acids Res 46: W329–W337.

Mészáros B, Simon I, Dosztányi Z. 2009. Prediction of protein binding regions in disordered proteins. PLoS Comput Biol 5: e1000376.

Namy O, Duchateau-Nguyen G, Hatin I, Hermann-Le Denmat S, Termier M, Rousset J. 2003. Identification of stop codon readthrough genes in Saccharomyces cerevisiae. Nucleic Acids Res 31: 2289–2296.

Neme R, Tautz D. 2013. Phylogenetic patterns of emergence of new genes support a model of frequent de novo evolution. BMC Genomics 14: 1–13.

Nepomnyachiy S, Ben-Tal N, Kolodny R. 2017. Complex evolutionary footprints revealed in an analysis of reused protein segments of diverse lengths. Proc Natl Acad Sci 114: 11703–11708.

Nielly-Thibault L, Landry CR. 2019. Differences between the raw material and the products of de novo gene birth can result from mutational biases. Genetics 212: 1353–1366.

Palmieri N, Kosiol C, Schlötterer C. 2014. The life cycle of Drosophila orphan genes. elife 3: e01311.

Papaleo E, Saladino G, Lambrughi M, Lindorff-Larsen K, Gervasio FL, Nussinov R. 2016. The role of protein loops and linkers in conformational dynamics and allostery. Chem Rev 116: 6391–6423.

Papandreou N, Berezovsky IN, Lopes A, Eliopoulos E, Chomilier J. 2004. Universal positions in globular proteins: From observation to simulation. Eur J Biochem 271: 4762–4768.

Postic G, Ghouzam Y, Chebrek R, Gelly J-C. 2017. An ambiguity principle for assigning protein structural domains. Sci Adv 3: e1600552.

Prabakaran S, Hemberg M, Chauhan R, Winter D, Tweedie-Cullen RY, Dittrich C, Hong E, Gunawardena J, Steen H, Kreiman G. 2014. Quantitative profiling of peptides from RNAs classified as noncoding. Nat Commun 5: 1–10.

Qiu L, Pabit SA, Roitberg AE, Hagen SJ. 2002. Smaller and faster: The 20-residue Trp-cage protein folds in 4 μs. J Am Chem Soc 124: 12952–12953.

Radhakrishnan A, Chen Y-H, Martin S, Alhusaini N, Green R, Coller J. 2016. The DEAD-box protein Dhh1p couples mRNA decay and translation by monitoring codon optimality. Cell 167: 122–132.

Ranwez V, Douzery EJ, Cambon C, Chantret N, Delsuc F. 2018. MACSE v2: toolkit for the alignment of coding sequences accounting for frameshifts and stop codons. Mol Biol Evol 35: 2582–2584.

Ranwez V, Harispe S, Delsuc F, Douzery EJ. 2011. MACSE: Multiple Alignment of Coding SEquences accounting for frameshifts and stop codons. PloS One 6: e22594.

Requião RD, Fernandes L, de Souza HJA, Rossetto S, Domitrovic T, Palhano FL. 2017. Protein charge distribution in proteomes and its impact on translation. PLoS Comput Biol 13: e1005549.

Rousseau F, Schymkowitz J, Serrano L. 2006a. Protein aggregation and amyloidosis: confusion of the kinds? Curr Opin Struct Biol 16: 118–126.

Rousseau F, Serrano L, Schymkowitz JW. 2006b. How evolutionary pressure against protein aggregation shaped chaperone specificity. J Mol Biol 355: 1037–1047.

Ruiz-Orera J, Messeguer X, Subirana JA, Alba MM. 2014. Long non-coding RNAs as a source of new peptides. elife 3: e03523.

Ruiz-Orera J, Verdaguer-Grau P, Villanueva-Cañas J, Messeguer X, Albà MM. 2018. Translation of neutrally evolving peptides provides a basis for de novo gene evolution. Nat. Ecol. Evol. 2: 890–896.

Scannell D, Zill O, Rokas A, Payen C, Dunham M, Eisen M, Rine J, Johnston M, Hittinger C. 2011. The Awesome Power of Yeast Evolutionary Genetics: New Genome Sequences and Strain Resources for the Saccharomyces sensu stricto Genus. G3 (Bethesda). Genet Soc Am. 2011; 1 (1): 11–25.

Schavemaker PE, Śmigiel WM, Poolman B. 2017. Ribosome surface properties may impose limits on the nature of the cytoplasmic proteome. Elife 6: e30084.

Schlötterer C. 2015. Genes from scratch–the evolutionary fate of de novo genes. Trends Genet 31: 215–219.

Schmitz JF, Ullrich KK, Bornberg-Bauer E. 2018. Incipient de novo genes can evolve from frozen accidents that escaped rapid transcript turnover. Nat Ecol Evol 2: 1626–1632.

Slavoff SA, Mitchell AJ, Schwaid AG, Cabili MN, Ma J, Levin JZ, Karger AD, Budnik BA, Rinn JL, Saghatelian A. 2013. Peptidomic discovery of short open reading frame– encoded peptides in human cells. Nat Chem Biol 9: 59.

Talmud D, Bresler S. 1944. On the nature of globular proteins. Comptes Rendus Dokady Acdémie Sci URSS 43 310–349.

Tautz D, Domazet-Lošo T. 2011. The evolutionary origin of orphan genes. Nat Rev Genet 12: 692–702.

Tendulkar AV, Joshi AA, Sohoni MA, Wangikar PP. 2004. Clustering of protein structural fragments reveals modular building block approach of nature. J Mol Biol 338: 611–629.

Thiaville PC, Legendre R, Rojas-Benítez D, Baudin-Baillieu A, Hatin I, Chalancon G, Glavic A, Namy O, de Crécy-Lagard V. 2016. Global translational impacts of the loss of the tRNA modification t6A in yeast. Microb Cell 3: 29.

Tretyachenko V, Vymětal J, Bednárová L, Kopecký V, Hofbauerová K, Jindrová H, Hubálek M, Souček R, Konvalinka J, Vondrášek J. 2017. Random protein sequences can form defined secondary structures and are well-tolerated in vivo. Sci Rep 7: 1–9.

Tusnády GE, Dosztányi Z, Simon I. 2005. PDB_TM: selection and membrane localization of transmembrane proteins in the protein data bank. Nucleic Acids Res 33: D275–D278.

Tusnády GE, Dosztányi Z, Simon I. 2004. Transmembrane proteins in the Protein Data Bank: identification and classification. Bioinformatics 20: 2964–2972.

UniProt Consortium. 2019. UniProt: a worldwide hub of protein knowledge. Nucleic Acids Res 47: D506–D515.

Vakirlis N, Acar O, Hsu B, Coelho NC, Van Oss SB, Wacholder A, Medetgul-Ernar K, Bowman RW, Hines CP, Iannotta J. 2020. De novo emergence of adaptive membrane proteins from thymine-rich genomic sequences. Nat Commun 11: 1–18.

Vakirlis N, Hebert AS, Opulente DA, Achaz G, Hittinger CT, Fischer G, Coon JJ, Lafontaine I. 2018. A molecular portrait of de novo genes in yeasts. Mol Biol Evol 35: 631–645.

Vakirlis N, McLysaght A. 2019. Computational prediction of de novo emerged protein-coding genes. In Computational Methods in Protein Evolution, pp. 63–81, Springer.

Wang M, Weiss M, Simonovic M, Haertinger G, Schrimpf SP, Hengartner MO, von Mering C. 2012. PaxDb, a database of protein abundance averages across all three domains of life. Mol Cell Proteomics 11: 492–500.

Wilson BA, Masel J. 2011. Putatively noncoding transcripts show extensive association with ribosomes. Genome Biol Evol 3: 1245–1252.

Wolf YI, Novichkov PS, Karev GP, Koonin EV, Lipman DJ. 2009. The universal distribution of evolutionary rates of genes and distinct characteristics of eukaryotic genes of different apparent ages. Proc Natl Acad Sci 106: 7273–7280.

Wu B, Knudson A. 2018. Tracing the de novo origin of protein-coding genes in yeast. MBio 9.

Yue J-X, Li J, Aigrain L, Hallin J, Persson K, Oliver K, Bergström A, Coupland P, Warringer J, Lagomarsino MC. 2017. Contrasting evolutionary genome dynamics between domesticated and wild yeasts. Nat Genet 49: 913–924.

Zhao L, Saelao P, Jones CD, Begun DJ. 2014. Origin and spread of de novo genes in Drosophila melanogaster populations. Science 343: 769–772.

